# Identification of age-associated proteins and functional alterations in human primary retinal pigment epithelium cells

**DOI:** 10.1101/2021.10.17.464744

**Authors:** Xiuxiu Jin, Jingyang Liu, Weiping Wang, Jiangfeng Li, Guangming Liu, Ruiqi Qiu, Mingzhu Yang, Meng Liu, Lin Yang, Xiaofeng Du, Bo Lei

## Abstract

**Background:** Retinal pigmented epithelium (RPE) has essential functions to nourish and support the neural retina, and is of vital importance in the pathogenesis of age-related retinal degeneration. However, the exact molecular changes of RPE in aging remain poorly defined.

**Methods:** We isolated human primary RPE (hRPE) cells from 18 eye donors distributed over a wide age range (10 - 67 years). A quantitative proteomic analysis was performed to analyze their intracellular and secreted protein changes, and potential age-associtated mechanisms were validated by ARPE-19 and hRPE cells.

**Results:** Age-stage related subtypes and age-associtated proteins and functional alterations were revealed. Proteomic data and verifications showed that RNF123 and RNF149 related ubiquitin-mediated proteolysis might be an important clearance mechanism in elimination of oxidative damaged proteins in aged hRPE. In older hRPE cells, apoptotic signaling related pathways were up-regulated and endoplasmic reticulum organization was down-regulated both in intracellular and secreted proteome.

**Conclusions:** This work paints a detailed molecular picture of human RPE in aging process and provides new insights for molecular characteristics of RPE in aging and related clinical retinal conditions.

## 1 BACKGROUND

Retinal degeneration (RD) is caused by gradual destruction of retinal cells ^1^, among which age-related macular degeneration (AMD) has become a leading cause of irreversible vision loss in the elderly worldwide ^2^. It is predicted that RD may progress concurrently with the aging of the population ^3^. The retina consists of the multi-layered neural retina and the monolayer of retinal pigment epithelium (RPE). RPE locates between the photoreceptors (RPs) and the vascular choroid, forming a critical barrier between the retina and the systemic circulation. Therefore, RPE performs many essential functions to nourish and support the neural retina, including recycling of vitamin A, daily phagocytosis of photoreceptor outer segment tips, and maintaining blood-retinal barrier. Evidence suggests that dysfunction and atrophy of RPE precedes many RDs, leading to the impairment of the neuroretina, resulting in significant vision loss ^4^.

The pathogenesis of age-related RD is complex and multifactorial, and the corresponding research models are limited. Oxidative stress is believed to contribute to the pathogenesis of age-related RD ^5^. Therefore, ample reports using oxidative damage agents, like paraquat and hydrogen peroxide (H_2_O_2_), to analysis the oxidative stress response mechanisms of RPE ^6-8^. Immortal cell lines such as ARPE-19 were commonly used to study human retina. However, these cell lines can differ significantly in some features relative to human primary RPE (hRPE) cells ^9^. Induced pluripotent stem cell (iPSC)-derived and human embryonic stem cell (hESC)-derived RPE cell lines may be promising models to study RPE. They have been proved to have behaviors similar to *in vivo* RPE and are currently being tested in clinical trials for RPE replacement ^10,11^. Several studies using induced iPSC-derived RPE cells from control and AMD patients to analyze the pathogenesis of AMD ^12,13^. However, unlike damaged RPE in AMD patients, iPSC-derived RPE cells from those patients are in a relative healthy status. hRPE cells are the ideal tool for investigating studies of RPE aging, but shortage of suitable donors and poor redifferentiation capacity limits their applications. Nevertheless, how hRPE changes in aging process is still unknown, which is a key factor to the pathogenesis of age-related RD. To speak of, RPE is mainly responsible for the secretion of mediators involved in the functional integrity of the RPs and the vascular choroid. Dysregulation of the secreted proteins in RPE was reported to be involved in the development of age-related RDs, especially in the loss of vascular invasion and barrier function ^14^.

In this study, we explored the intracellular and secreted protein changes of the hRPE cells, obtained from donors distributed over a wide age range. A quantitative mass spectrometry (MS)-based proteome analysis (tandem mass tag, TMT) was performed to characterized those overrepresented and underrepresented proteins with age. These data was made inferences about molecular pathways affected by aging in hRPE cells. This work provided a rich resource to study the impact of aging on hRPE, and presented solid data for future research on RPE and related clinical retinal conditions.

## 2 MATERIALS AND METHODS

### 2.1 Cell Culture

Human primary RPE (hRPE) cells were obtained from eyes of donors (Henan Eye Bank) according to previously published protocols ^15^. Briefly, eyes were cut circumferentially above the equator, then the lens, vitreous, and iris tissue were removed. Choroid was separated from the sclera. hRPE cells were removed from choroid by incubation in 7.5% trypsin-EDTA solution. ARPE-19 cells were purchased from American Type Culture Collection (ATCC CRL-2302, Manassas, VA, USA).

Both hRPE and ARPE-19 cells were grown in a mixture (1:1) of Dulbecco’s Modified Eagle’s Medium (DMEM): Nutrient F-12 Ham (Gibco, #C11995-065) supplemented with 10% fetal bovine serum (FBS, NATOCOR, #SFBE), 100 U/mL penicillin and 100 U/mL streptomycin (Gibco, #15140-122) and were cultured in 37 °C with 5% CO_2_ incubator.

### 2.2 Compounds and Antibodies

The antibodies and compounds used were commercially obtained: anti-RPE65 antibody (Abcam, #ab13826, 1:2000), anti-RNF123 antibody (BOSTER, #A09642-1, 1:500), anti-RNF149 (Bioss, #bs-9228R, 1:500), anti-ubiquitin antibody (Cell Signaling Technology, #3933, 1:2000), anti-β-Actin (Huabio, #EM21002, 1:5000), H_2_O_2_ (Sigma-Aldrich, #323381), 40,6-diamidino-2-phenylindole (DAPI, Sigma-Aldrich, #9542), lutein (Sigma-Aldrich, #07168).

### 2.3 Cell Viability Assay

The effects of H_2_O_2_ on the viability of hRPE cells with different ages were determined using cell counting kit-8 (CCK-8, Sigma-Aldrich, #96992) according to the manufacturer’s instructions. Briefly, approximately 4000 hRPE cells were seeded in 96-well plates and allowed to adhere for 12 h, then gradient concentrations of H_2_O_2_ (supplement with 0 μM, 5 μM or 10 μM lutein) were added and incubated with the cells for another 48 h before WST-8 was added to each well. After incubation for another 1h, cells were measured using a multifunction microplate reader (PerkinElmer, Waltham, MA, USA). Cells cultured without treatment were used as control.

### 2.4 Immunofluorescence Staining

hRPE and ARPE-19 cells were seeded in 12-well plates and cultured to 80% confluence, fixed with 4% paraformaldehyde for 30 minutes at room temperature, followed by three times wash with PBS (pH = 7.4), permeabilized with 0.2% Triton X-100 for 20 minutes, then washed with PBS three times. The cells were then blocked in PBS containing 5% BSA for 2 hour and incubated with anti-RPE65 antibody (Abcam, #ab13826, 1:1000) overnight at 4°C. Cells were washed three times in PBS then incubated with fluorescent-conjugated secondary antibody (Cell Signaling Technology, 1:250, #4408) for 1.5 h at room temperature, followed by 5 minutes of nuclear staining with DAPI. All fluorescent images were taken with a fluorescence microscope (OLYMPUS IX73, Tokyo, Japan).

### 2.5 Intracellular and Secreted Protein Lysate Preparation

For secreted proteins, hRPE cells were washed 3 times with PBS (pH = 7.4) before adding 5 mL of serum-free RPE media. Cells were cultured for 24 h and then the media was collected, and the secretome was washed and concentrated by a centrifugal ultrafiltration membrane (Amicon Ultra-15, MWCO 15 KDa). For intracellular proteins, hRPE cells were washed twice with ice-cold PBS, harvested with a cell scraper and collected by low-speed centrifugation. Then both of the secretome (for secreted proteins) and hRPE cells (for intracellular proteins) were lysed with RIPA buffer containing protease inhibitor coctail (1% NP-40, 0.5% (w/v) sodium deoxycholate, 150 mM NaCl, 50 mM Tris (pH = 7.5)), followed by 3 min sonication under the condition of 3 s on and 10 s off with 195 watt of JY92-IIN (NingBoXinYi, China). Protein quantification was performed by Bradford (Bio-Rad, #5000205).

### 2.6 Proteomics Sample Preparation

Each protein sample was diluted to 1 μg/μL (50 μg for intracellular protein analysis and 20 μg for secreted protein analysis) by 100 mM tetraethylammonium bromide (TEAB, Sigma-Aldrich, #15715-58-9), reduced by Tris (2-carboxyethyl) phosphine (TCEP, Sigma-Aldrich, #C4706) with a final concentration of 10 mM at 56 °C for 1 h, alkylated with iodoacetamide (IAA, Sigma-Aldrich, #I1149) with a final concentration of 20 mM in the dark at room temperature for 30 min, precipitated with methanol, chloroform and water (CH_3_OH:CHCl_3_:H_2_O = 4:1:3), followed by overnight digestion with trypsin (Progema, #V5117, 1:50 enzyme to protein). The tryptic peptides of each sample were labeled with TMT-10 plex (Thermo Fisher Scientific, #90113CH) reagents according to the manufacturer’s protocol. One batch of 10-plex TMT kits was used to label 9 intracellular (or secreted) samples and one mixture of hRPE intracellular proteins (or secreted proteins). After quenching with 5% hydroxylamine, TMT-labeled peptides of each batch of TMT-10 plex were mixed and desalted by C18 column. Then the desalted peptides were fractionated by Agilent Poroshell HPH C18 column (250 × 4.6 mm, OD 4 μm) on Agilent 1260 instrument.

Buffer A (2% ACN, 10 mM NH_4_COOH, pH = 10) and a nonlinear increasing concentration of buffer B (90% ACN, 10 mM NH_4_COOH, pH = 10) were used for peptide separation at a flow rate of 1 mL/min. A standard 120 min gradient was used as follows, 0%-8% B in 10 min; 8%-35% B in 70 min; 35%-60% B in 15min; 60%-70% B in 10 min; 70%-100% B in 15 min. The peptides mixture was separated into 120 fractions and combined by a concatenation strategy into 24 fractions (1&25&49&73&97, …, 24&48&72&96&120). The combined fractions were used for further LC-MS/MS analysis.

### 2.7 LC-MS/MS analyses

Peptides were dissolved in buffer A (0.1% formic acid, FA) and then directly loaded onto a reversed-phase analytical column (Acclaim PepMap RSLC, Thermo). The gradient was comprised of an increase from 6% to 22% solvent B (0.1% FA in 98% CH_3_CN) over 42 min, 22% to 30% in 12 min and climbing to 80% in 3 min then holding at 80% for the last 3 min, all at a constant flow rate of 500 nL/min on an EASY-nLC 1000 Ultra Performance Liquid Chromatography (UPLC) system. The peptides were then subjected to nanospray ionization (NSI) source followed by tandem mass spectrometry (MS/MS) in Orbitrap Fusion Lumos™ instrument (Thermo Scientific) coupled online to the UPLC. Intact peptides were detected in the Orbitrap at a resolution of 60,000. Peptides were selected for MS/MS using normalized collision energy (NCE) setting as 32 and ion fragments were detected in the Orbitrap at a resolution of 15,000. A data-dependent procedure that alternated between one MS scan followed by 20 MS/MS scans was applied for the top 20 precursor ions above a threshold intensity greater than 1e4 in the MS survey scan with 30.0 s dynamic exclusion. The electrospray voltage applied was 2.4 kV. Automatic gain control (AGC) was used to prevent overfilling of the orbitrap. 5e4 ions were accumulated for generation of MS/MS spectra. For MS scans, the m/z scan range was 350 to 1550. Fixed first mass was set as 100 m/z.

### 2.8 MS Data Processing

The resulting MS/MS data was processed using MaxQuant with integrated Andromeda search engine (version 1.5.2.8). Tandem mass spectra were searched against the human UniProt database (20239 entries, 2017/09) concatenated with reverse decoy database. Trypsin/P was specified as cleavage enzyme allowing up to 2 missing cleavages. Mass error was set to 10 ppm for precursor ions and 0.02 Da for fragment ions. Carbamidomethylation on cysteine was specified as fixed modification, and oxidation on methionine and acetylation on protein N-terminal were specified as variable modifications. False discovery rate (FDR) thresholds for protein, peptide and modification site were specified at 1%. Minimum peptide length was set at 7. For quantification method, TMT-10plex was selected. All the other parameters in MaxQuant were set to default values. Intensities were extracted and normalized based on their total intensity and the internal standard to correct the sample loading difference. The zero intensity proteins, reverse and contaminant were excluded.

### 2.9 Proteomics informatics

Pathway analysis was performed using the R package XGR ^16^ using the GOBP and Reactome databases. GSEA analysis was conducted using tumor hallmark database (Version: h.all.v 7.0.symbols.gmt [Hallmarks]) ^17^. Functional enrichment analysis of biological relevance of the hub genes and their regulators in proteins specifically changed in secretome was performed using Cytoscape (version 3.7.2) plugged with ClueGO (version 2.5.7) ^18^. Consensus clustering was performed using the R package ConsensusClusterPlus ^19^. Samples were clustered using euclidean distance as the distance measure.

### 2.10 Protein co-expression network analysis

Protein co-expression network analysis was performed with the R package (weighted gene co-expression network analysis (WGCNA)) using entire secreted proteomic data set of all identified proteins according to previous report ^20^. The eigengene value of each module was calculated to test the correlation with each sample ^21^. Before creating the network model, the distribution of the entire dataset was performed. The hub genes from the model were screened by calculating the connectivity degree of each gene with Cytoscape (version 3.7.2) ^22^.

### 2.11 Western blot analysis

Protein extracts was separated on 10% sodium dodecyl sulfate-polyacrylamide gel electrophoresis, and then transferred onto a polyvinylidene difluoride (PVDF) membrane (Millipore, Burlington, MA, USA). After blocking with 5% milk solution in TBST (Tris buffered saline with Tween) for 1 h, membranes were incubated with 3% milk containing appropriate primary antibodies overnight at 4 °C, followed by 2 h of incubation with horseradish peroxidase conjugated secondary antibodies (Cell Signaling Technology, 1:5000). Signals of target protein bands were detected using Chemiluminescent detection reagent.

## 3 RESULTS

### 3.1 Workflow and General Overview

An overview of the experimental strategy is shown in Figure 1. Eyes from male donors within 72 h after death were obtained from Henan Eye Bank (Zhengzhou, China). Previous medical and ocular histories were assessed to exclude donors with any eye disease. Then the eyes were divided into three groups according to the ages of the donors, including the young (Y, 10 - 18 y, n = 4), the middle-aged (M, 35 - 41 y, n = 7) and the old (O, 55 - 67 y, n = 7) (Supplementary Table 1). hRPE cells were isolated immediately and cultured for 2 passages. To confirm hRPE cell line, immunofluorescence staining assay was performed by RPE-specific antibody, anti-human RPE65 ^23^. RPE65 is a cytosolic protein, which locates in the cytoplasm around the 4’,6-diamidino-2-phenylindole (DAPI)-positive nucleus. Clearly, both hRPE and ARPE-19 cells displayed similar morphology and RPE65 staining results (Supplementary Figure 1A), which indicated that those cell lines we collected from eyes of the donors were RPE-derived cell lines.

**Figure 1.**
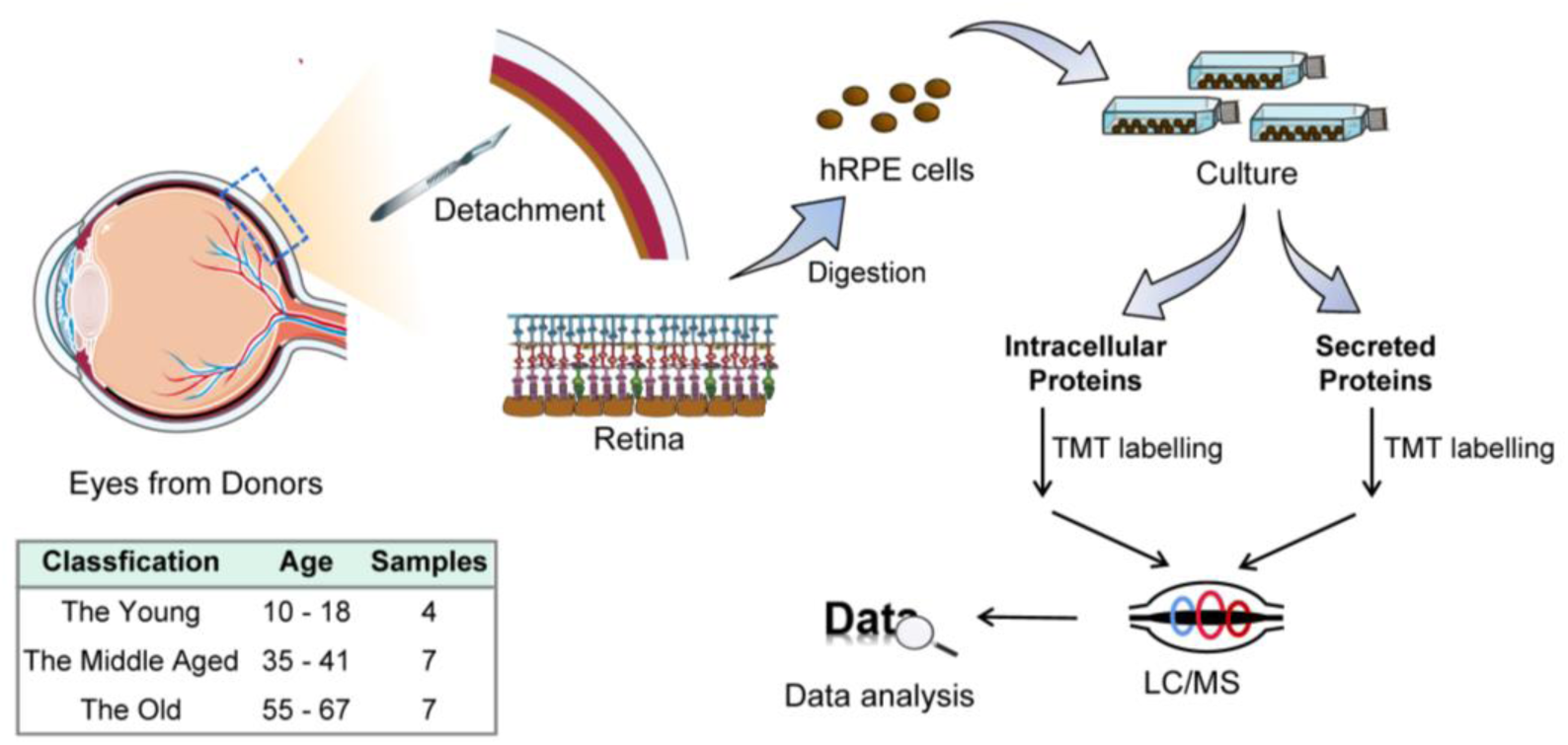
Graphical illustration of the workflow in this study. hRPE cells were isolated from eyes of the donors, after two passages, intracellular and secreted proteins of hRPE cells were extracted and analyzed by TMT^10^-based proteomic profiling.

To obtain the changing patterns of intracellular and secreted proteome between the hRPE cell lines with different ages, we performed quantitative proteomics using TMT-10plex isobaric labeling reagents. At the third passage of hRPE cells, intracellular and secreted proteins were extracted and digested for further TMT labelling. To eliminate batch effects, samples were randomly labeled by TMT-10plex (Supplementary Figure 1B). Mixture of intracellular (or secreted) protein samples was labeled as internal stand in each batch of TMT-10plex, and further analyzed by liquid chromatography tandem-MS (LC-MS) analysis. A between-sample normalization by the median of intensities was performed before any quantification analysis (Supplementary Figure 1C & D). Those overrepresented and underrepresented proteins in older donors were made inferences about molecular pathways affected by aging in hRPE cells.

### 3.2 Proteomic Features of hRPE cells

In intracellular samples, we identified 74,502 unique peptides from 7921 proteins with average sequence coverage of 25.1% (protein false discovery rate [FDR] <1%). And 27,930 unique peptides from 3843 secreted proteins with average sequence coverage of 17.4% (FDR <1%) were identified (Figure 2A, Supplementary Data 1). Among those identified proteins, 5735 intracellular proteins and 2455 secreted proteins were quantified in all samples (n=18). Quality controls including number of quantified proteins and coefficient of variation (CV) of the Y, the M and the O groups were further analyzed. Results showed that number of quantified proteins and the median CV values of the Y, the M and the O groups were comparable (Supplementary Figure 2A-D). And the CV values were no more than 2.55% in intracellular samples, and not exceed 3.22% in secreted samples. The top 10 most abundant intracellular proteins accounts for 6.26% of the total spectral abundance, including PLEC, AHNAK, MYH9, FLNA, VIM, FLNC, PRKDC, ACTG1, MACF1, DYNC1N1 (Figure 2B). Besides, COL1A1, COL1A2, C3, FN1, FLNA, ACTN4, FLNC, FBN1, AGRN and LAMB1 are the top 10 most abundant secreted proteins, which accounted for 7.69% (Figure 2C). The above data indicated that the protein quantification across the 18 TMT experiments was robust.

**Figure 2.**
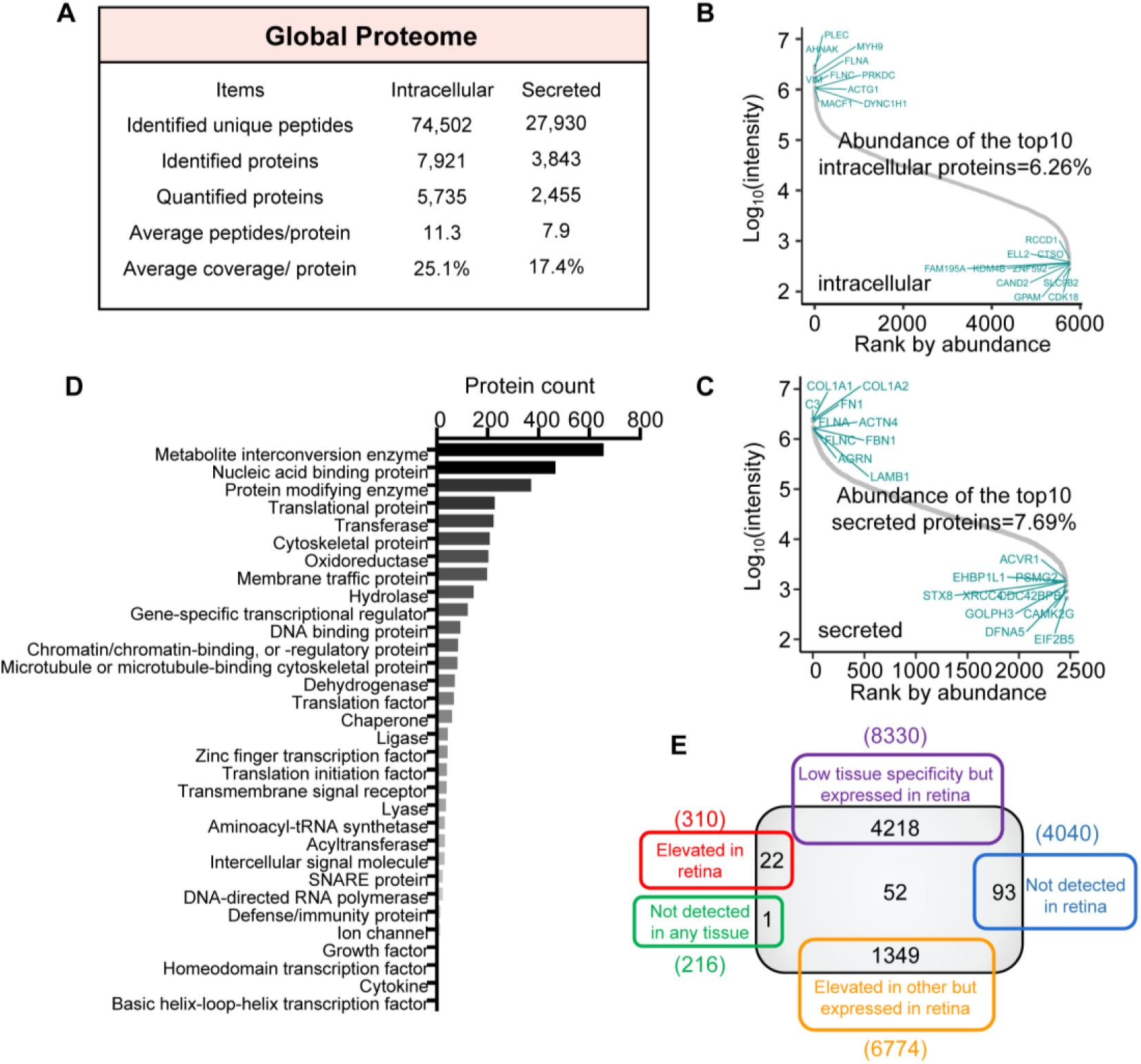
Proteome overview of hRPE cells. (A) Global proteome of hRPE cells. (B-C) Protein abundance analysis of the quantified intracellular (B) and secreted (C) proteins. The x-axis indicates protein ranking orders according to their abundance. The y-axis represents the log_10_ (intensity) of each protein. The top 10 most abundant proteins are labeled in the top left. (D) Number representations of indicated PANTHER protein categories of the quantified intracellular proteins. (E) Venn diagram of the numbers of expressed genes in retina on the mRNA level (The Human Protein Atlas Database-Tissue Atlas (Retina)) and the quantified intracellular proteins in hRPE cells. Grey box represents intracellular quantified proteins in this study.

To further characterize the properties of hRPE, we performed PANTHER protein class (version 15.0) and Gene Ontology (GO) subcellular location analysis. PANTHER protein class analysis indicated that those quantified intracellular proteins covered a broad range of protein classes, e.g., metabolite interconversion enzyme (655 proteins), nucleic acid binding protein (465 proteins) and protein modifying enzyme (369 proteins) (Figure 2D, Supplementary Data 1). Subcellular location analysis showed that most of the intracellular proteins belonged to membrane (3157 proteins), nucleus (2895 proteins) and vesicle (1777 proteins) (Supplementary Figure 2E, Supplement Data 1). RPE cells are pigment-rich cells with obvious secretory properties ^24^, and we also quantified 447 secretory vesicle proteins and 81 pigment granule proteins.

Subsequently, we matched intracellular proteins with the expressed genes in retina on mRNA level (The Human Protein Atlas Database-Tissue Atlas (Retina)). Venn diagram showed that 22 of the quantified proteins were reported to be elevated in retina than other tissues. Besides, 93 proteins not detected in retina and one protein not detected in any tissue were also quantified in our research (Figure 2E, Supplement Data 1). Next, those quantified secreted proteins were matched with the expressed genes of secreted proteins and vesicles on mRNA level, as predicted in The Human Protein Atlas Database-The Cell Atlas. Venn diagram showed that 725 quantified proteins were predicted as secreted proteins, and 302 proteins belong to vesicles (Supplementary Figure 2F, Supplementary Data 1). Therefore, our results provided broad coverage of the proteins of hRPE in individuals.

### 3.3 The Proteomic Variances of hRPE in Aging Process

To illustrate the relationship between age-stages and the related changes of molecular functions in hRPE, we employed consensus clustering to identify age-stage related subtypes (Figure 3A, Supplement Data 2). In total, 10 clusters (C1 - 10) were enriched with significant changes (p < 0.05) in molecular function. Apparently, those pathways enriched by proteins in C1, C5 and C6 are relatively stable in Y and M stages, but changes significantly in O. When entering O stage, C1 related pathways such as intra golgi vesicle mediated transport, Copii coated vesicle budding and golgi vesicle budding were up-regulated. C5 and C6 related pathways including neuroinflammatory response, positive regulation of synapse assembly, *etc*., were down-regulated in O. As indicated by C7, proteins enriched in long chain fatty acyl CoA metabolic process were decreased in aging process. Therefore, pathways enriched by proteins in C1, C5, C6 and C7 may be closely related to the pathological process of age-related RD. However, comparing with O and Y, proteins in M stage exhibited down-regulated expression levels in pathways enriched by proteins in C2, C3 and C8, including cellular response to glucose starvation (C2), response to misfolded protein (C3) and regulation of chondrocyte differentiation (C8). Besides, proteins in M stage showed specific up-regulation in C9 and C10, such as macrophage activation (C9) and negative regulation of leukocyte proliferation (C10).

**Figure 3.**
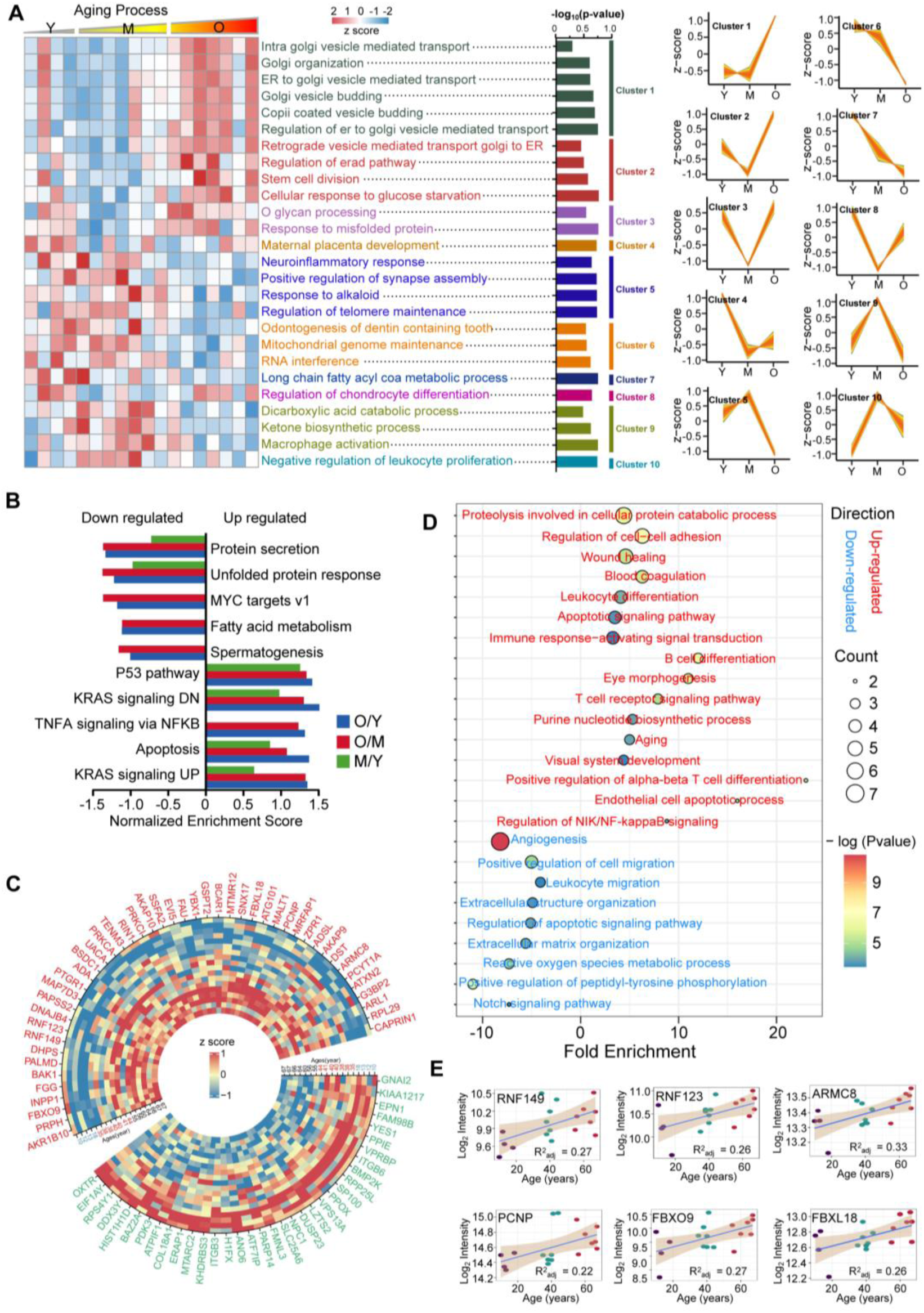
Visualization of intracellular changed proteins and their biological pathways. (A) Significant discrete clusters to illustrate the relative expression changes of the intracellular proteomics. Heatmap of each sample (n=18) for their subtype associated molecular functions are shown on the left. Significant clusters shown on the right are clustered by mFuzz. (B) GSEA analysis of the quantified proteins using hallmark database (version 7.0). GSEA was performed using version 4.0.3 of the GSEA desktop application. (C) Heatmap of the age-associated proteins in hRPE cells. Age-associated proteins were defined as R > 0.5 (or < -0.5, protein intensities *vs* ages), and P < 0.05 (Y *vs* O). (D) Biological process enrichment using GO database with the age-associated proteins. (E) Age-dependent increase of proteins enriched in proteolysis involved in cellular protein catabolic process (RNF149, RNF123, ARMC8, PCNP, FBXO9 and FBXL18). Simple linear regression was shown for age (x-axis) and protein intensity (y-axis) correlation.

To further analyze the relationships between different stages (O *vs* Y, M *vs* Y and O *vs* M) and reveal the influences of aging process on hRPE, proteomic data were analyzed by gene set enrichment analysis (GSEA) (hallmark database, version 7.0). It was found that O/Y and O/M exhibited similar changing patterns in most enriching hallmarks (Figure 3B, Supplement Data 2). This phenomenon suggested these proteins under wend noticeable changes when approaching O stage, which might be the reason that RPE-related diseases were prone to morbidity in old age ^3^. Besides, in aging process, hRPE cells was down-regulated in proteins secretion, unfolded protein response, *etc*., and up-regulated in P53 pathway, apoptosis, *etc*.. Above hallmark changes may contribute to the pathogenesis of RPE aging.

### 3.4 Age-associated proteins and the related functions

To disclose age-associated proteins in hRPE cells, we calculated the Pearson’s correlation coefficient (R) of the intensities of the quantified proteins between different ages. Cutoff value of R was defined as 0.5, and P-value (Y *vs* O) was set to less than 0.05. Forty-six proteins (such as AKR1B10, PRPH, FBXO9, *etc*.) were determined as up-regulated proteins, and 35 proteins (such as OXTR, EIF1AY, RPS4Y1, *etc*.) were down-regulated proteins (Figure 3C, Supplement Data 3). Specifically, the up-regulated AKR1B10 was reported to exert a protective role through eliminating oxidative stresses and was identified as a biomarker in older hepatocellular carcinoma patients ^25, 26^.

Next, enrichment analysis was performed to reveal potential functional changes of hRPE affected by age-associated proteins. KEGG (Kyoto Encyclopedia of Genes and Genomes) pathway analysis revealed that RAP1 signaling pathway enriched by PRKCA, PRKCI, BCAR1, GNAI2 and ITGB3 might change in aging process (Supplementary Figure 3C, Supplementary Data 3). RAP1 signaling pathway was reported to protect telomeres of senescent cells from DNA damage ^27^, which indicated that these 5 proteins might contribute to DNA damage related aging process. GO analysis showed that down-regulated proteins including ITGB3, SP100, EPN1, ERAP1, COL18A1, FMNL3 and ATP5IF1 were enriched in angiogenesis (Figure 3D, Supplementary Data 3). Angiogenesis is a pathological hallmark in many retinal vascular diseases ^28^. Therefore, the 7 down-regulated proteins might associate with the pathogenesis of these diseases. In addition, 6 up-regulated proteins including RNF149, RNF123 ARMC8, FBXO9, PCNP and FBXL18 were enriched significantly in proteolysis involved in cellular protein catabolic process (Figure 3D-E, Supplementary Data 3). RNF149 and RNF123 are two E3 ubiquitin-protein ligases, which regulate protein degradation through ubiquitin-mediated proteolysis ^29^.

Ubiquitin-mediated proteolysis is critical in proteostasis, which counteracts the insults accumulated to proteins by removing damaged proteins during aging.

### 3.5 Ubiquitin-mediated proteolysis in aging process of hRPE

To demonstrate the reliability of our proteomic data and reveal the relationships between ubiquitin-mediated proteolysis and aging process, western blotting (WB) was performed. We found that RNF123, RNF149 and the ubiquitinated proteins were up-regulated with age (Figure 4A). Oxidative stress is a hallmark of aging and previous reports using H_2_O_2_-treated cells to analysis the mechanisms of aging ^30, 31^. To further verify the effect of RNF123 and RNF149 related protein ubiquitination in RPE aging process, ARPE-19 cells with gradient concentrations of H_2_O_2_ treatment was analyzed. WB results showed that RNF123, RNF149 and the ubiquitinated proteins were up-regulated with increasing concentrations of H_2_O_2_, this phenomenon can be reversed by lutein, a natural antioxidant (Figure 4B). Then, hRPE cells from Y and O donors were further treated by gradient concentrations of H_2_O_2_. CCK8 viability results showed that hRPE cells of O donors were more vulnerable to oxidative damage (H_2_O_2_) (Figure 4C). WB results indicated that oxidative damage was related with the up-regulation of ubiquitylation of proteins, which can be rescued by lutein as well (Figure 4D). Therefore, RNF123 and RNF149 related protein ubiquitination may be an important clearance mechanism in elimination of oxidative damaged proteins in aged hRPE.

**Figure 4.**
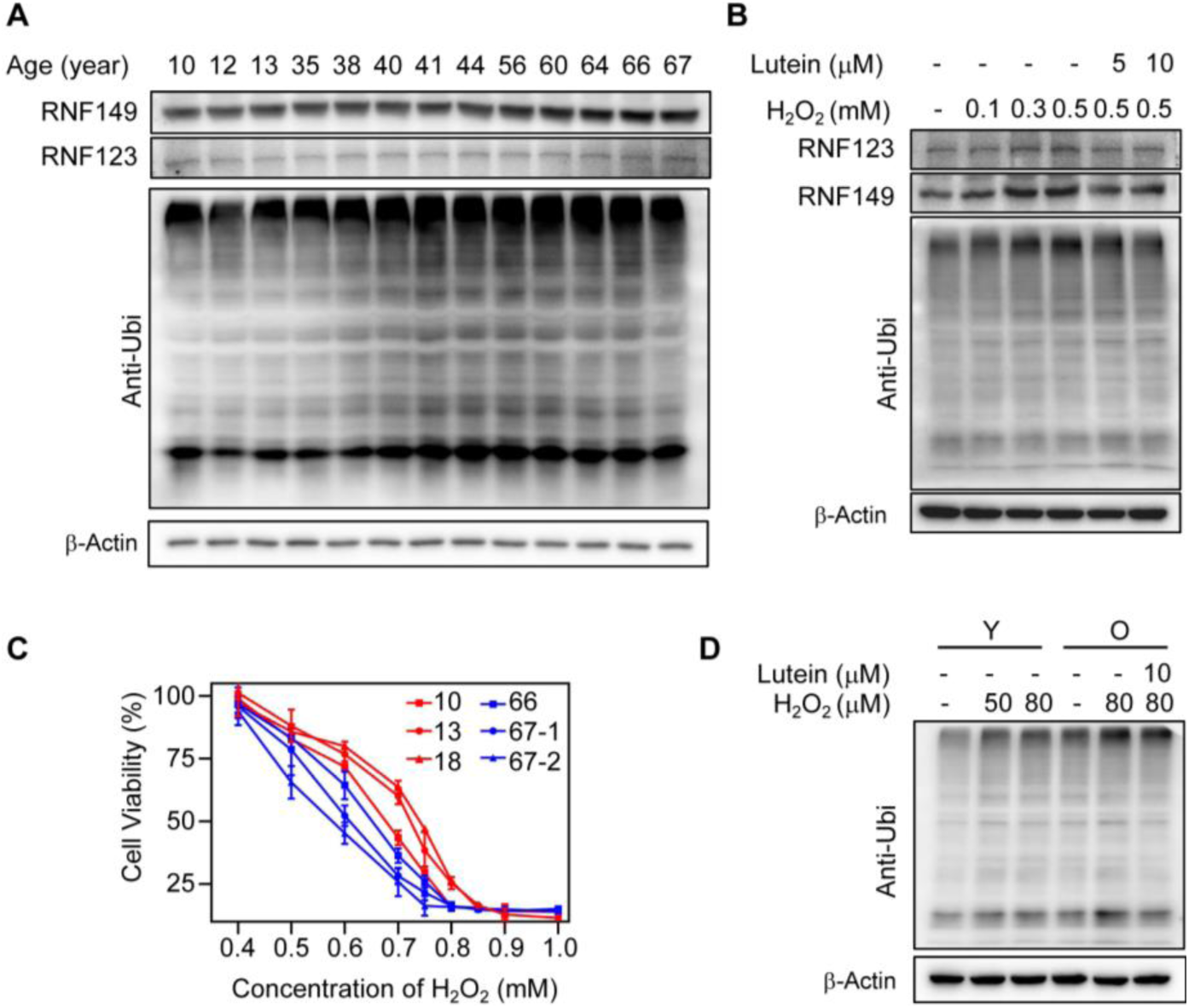
Ubiquitin-mediated proteolysis in aging process of hRPE. (A) Western blotting (WB) analysis of the hRPE cells with different ages. RNF123, RNF149 and the ubiquitinated proteins in hRPE were up-regulated with age. (B) WB analysis of the ARPE-19 cells, pretreated with lutein for 6 h before stimulation with gradient concentrations of H_2_O_2_ for 30 min. RNF123, RNF149 and the ubiquitinated proteins were up-regulated in ARPE-19 cells with increasing concentrations of H_2_O_2_, and this phenomenon can be reversed by lutein. (C) Cell viability of hRPE cells with different ages after H_2_O_2_stimulation. hRPE cells of O donors were more vulnerable to H_2_O_2_ stimulation. (D) WB analysis of the O and the Y hRPE cells confirms the H_2_O_2_ stimulation is related with the up-regulation of ubiquitylation of proteins. 67-year-old donor for the old, and 13-year-old donor for the young.

### 3.6 Age-associated secretory phenotype and proteins

Mediators secreted by RPE are involved in the functional integrity of the RPs and the vascular choroid ^14^. To reveal age-associated secretory phenotype, we employed consensus clustering analysis of the secreted proteins. In total, 9 clusters (C1 - 9) were enriched (Figure 3A, Supplementary Figure 4A, Supplement Data 4). Apparently, pathways enriched by proteins in C2 were up-regulated with age. These pathways were related with apoptotic signaling pathways, including regulation of intrinsic apoptotic signaling and intrinsic apoptotic signaling. The up-regulation of apoptotic signaling related pathways were consistent with intracellular GSEA analysis, for example the up-regulation of P53 pathway and apoptosis (Figure 3D). However, pathways like negative regulation of cellular component organization and ER organization enriched by proteins in C9 were down-regulated with age. ER is the site of entry of proteins into the secretory pathway ^32^. Therefore, the down-regulated ER organization in secretome was consistent the intracellular results that proteins in protein secretion and unfolded protein response were down-regulated in hRPE in aging process.

Age-associated secreted proteins were further analyzed by using p < 0.05 (Y *vs* O) and R > 0.5 (or R < -0.5, protein intensities between different ages). As shown in Figure 5B, 38 proteins (SRGN, DIS3, *etc*.) were determined as up-regulated, and 16 proteins (MYEF2, SPARCL1, *etc*.) were down-regulated. Many of the age-associated secreted proteins were reported to be associated with age-related diseases or as age-related proteins. For example, increased expression of SRGN was confirmed to be associated with greater aggressiveness in inflammation ^33,34^, and SPARCL1 was reported to be enriched in serum from young mice ^35^.

**Figure 5.**
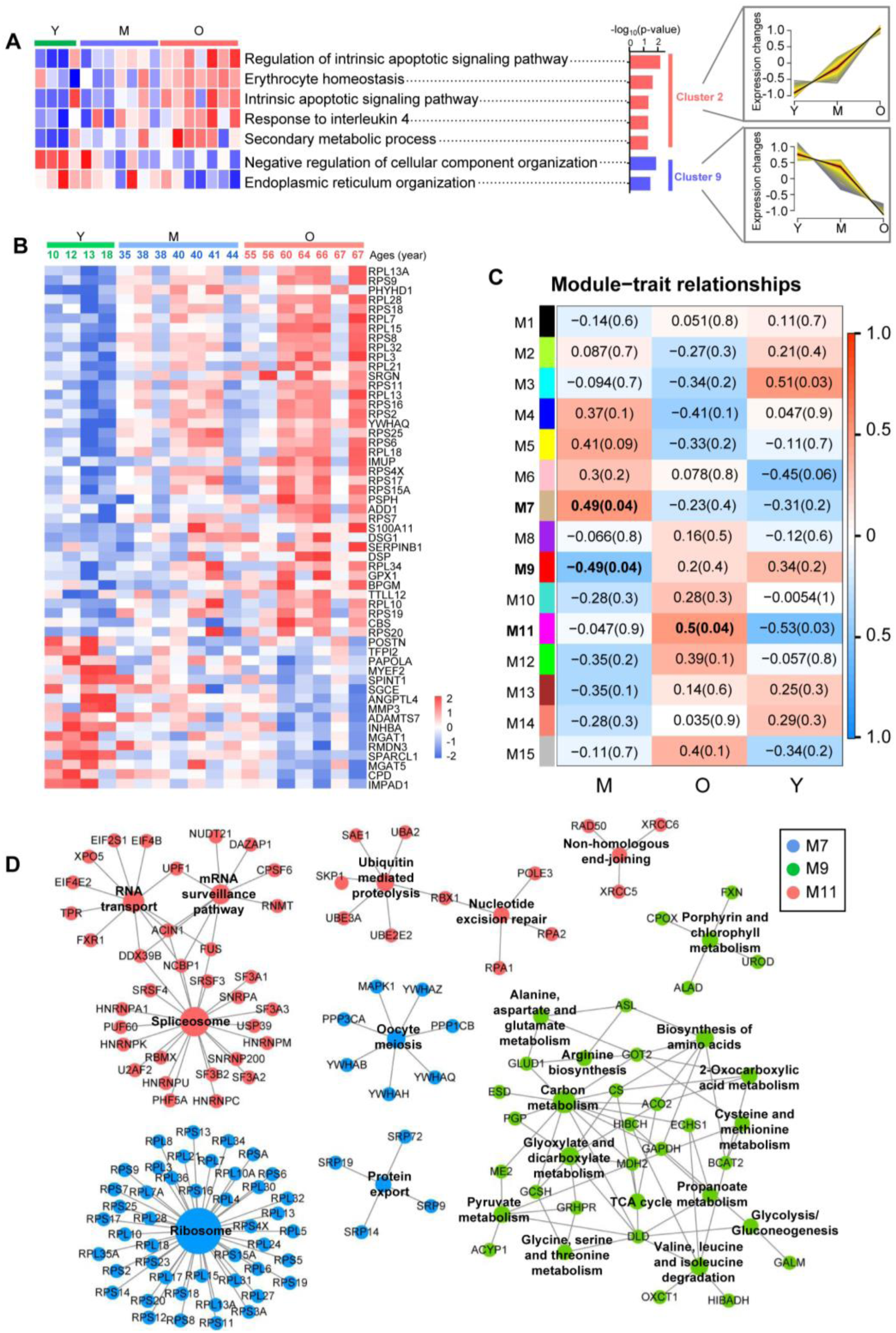
Visualization of age-associated secretory phenotype. (A) Significant down-regulated and up-regulated clusters to illustrate the relative expression changes of the secreted proteomics. Heatmap of each sample (n=18) for their subtype associated molecular functions are shown on the left. Clusters shown on the right are the up-regulated and down-regulated clusters which were clustered by mFuzz. (B) Heatmap of the age-associated secreted proteins. Age-associated secreted proteins were defined as R > 0.5 (or <-0.5, protein intensities *vs* ages), and P < 0.05 (Y *vs* O). (C) Identification of aging-relevant protein modules associated with age traits. Module-trait relationships were determined by biweight midcorrelation between module eigenprotein expression and the age stages. Correlation coefficients are indicated on the left with corresponding P-values in brackets on the right. Significant correlations (P < 0.05) are labeled in bold text. (D) KEGG network depiction of protein co-expression modules. Nodes represent proteins and edges (lines) indicate connections between the nodes. Blue represents M7 (positive correlations), Green represents M9 (negative correlations), and red represents M11 (positive correlations).

### 3.7 Co-expression network analysis uncovers secreted protein network alterations associated with aging

To get systems-level insights into age-associated secretome changes of hRPE, we performed protein co-expression network analysis by using WGCNA. In total, 15 network modules of strongly co-expressed proteins were identified (Figure 5C, Supplementary Data 4). These modules were color coded according to the convention of WGCNA, and then labeled by M1 to M15. To identify age-relevant modules, the module-trait relationships determined by biweight midcorrelations between each module eigenprotein (the module representative which summarizes protein expression profiles in the module) and sample variables were assessed. Three modules that were significantly correlated (p < 0.05) with age were identified, including two positive correlated modules (M7, M11) and one negatively correlated module (M9).

To gain insights into the biological roles of age-related modules, molecular and functional characteristics of these 3 modules were further analyzed based on KEGG and GO database. KEGG enrichment analysis revealed that positive correlated modules (M7, M11) were significantly enriched in RNA related pathways (including RNA transport, mRNA surveillance pathway, spliceosome), ubiquitin mediated proteolysis, nucleotide excision repair, non-homologous end-joining, ribosome, oocyte meiosis and protein export. Specifically, ubiquitin mediated proteolysis enriched by secreted proteins including SAE1, UBA2, RBX1, UBE2E2, UBE3A, SKP1, was also defined as an age-associated functional changes in intracellular proteomics analysis (Figure 3D), which was further verified by WB (Figure 4A). In addition, pathways related to small molecular biosynthesis or metabolism were enriched in negatively correlated module (M9), such as alanine, aspartate and glutamate metabolism, arginine biosynthesis, biosynthesis of amino acids *etc*.. GO enrichment analysis indicated that categories of structural constituent of ribosome (M7), translation regulator activity (M11), processes linked to the binding (M7 & M11, including ribonucleoprotein complex binding, cadherin binding, single-stranded DNA binding, cell adhesion molecule binding, rRNA binding, *etc*.) were up-regulated with age (Supplementary Figure 5B, Supplementary Data 4). However, enzymatic activity (including hydro-lyase activity, carbon-oxygen lyase activity, lyase activity, etc.), NAD binding and coenzyme binding were down-regulated with age.

### 3.8 Specifically Changed Secreted Proteins in hRPE Cells in Aging

Many proteins are present both inside the cell and within the extracellular space, and those changed specifically in secreted part may be essential for inter-cellular communication in aging process. Therefore, proteins quantified both in intracellular and secreted hRPE samples were further analyzed. Venn diagram showed that, 2067 proteins were shared by intracellular and secreted samples, and 147 of them were significantly changed (p < 0.05) in secreted proteins, including IMPAD1, CPD, RMDN3, MGAT1, MYEF2, PAPOLA, *etc*. (Figure 6A & B, Supplementary Data 5). Functional enrichment analysis of the specifically changed secreted proteins was performed by ClueGo software (version 2.5.7) in Cytoscape (version 3.7.2). Six clusters of 108 significantly enriched GO terms (Biological Processes) were identified, including SRP-dependent cotranslational protein targeting to membrane, myeloid cell homeostasis, regulation of ventricular cardiac muscle cell action potential, neutrophil degranulation and ribosome biogenesis, and alpha-amino acid biosynthetic process.

**Figure 6.**
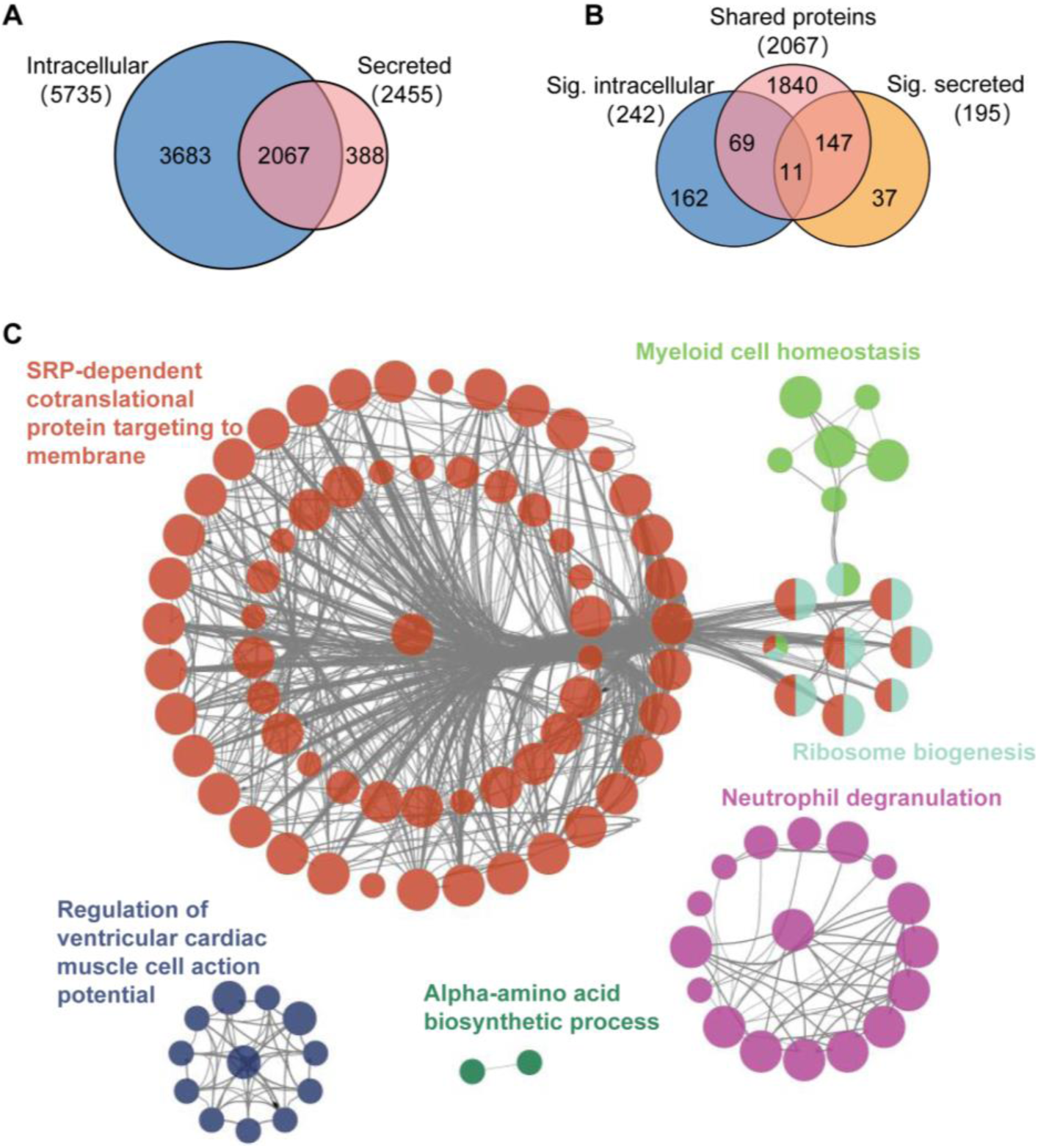
Visualization of the specifically changed secreted proteins. (A) Venn diagram shows the overlap between the quantified intracellular (blue) and secreted (red) proteins. (B) Venn diagram shows the overlap of significantly regulated intracellular (blue) and secreted (yellow) proteins versus the shared proteins of Venn diagram (A) (red). (C) Functional enrichment analysis of the specifically changed secreted proteins. Analysis was performed by ClueGo software (version 2.5.7) in Cytoscape (version 3.7.2).

These biological processes may play an important role in inter-cellular communication in RPE aging.

## 4 DISCUSSION

RPE performs many essential functions to nourish and support the neural retina, and is of vital importance in the pathogenesis of age-related RD, such as AMD ^36^.

However, the exact molecular changes of RPE in aging process that mediate dysfunction remain poorly defined. Ample reports using immortal cell lines (such as ARPE-19), iPSC-derived or hESC-derived RPE cell lines to explore the pathology of RPE-related disease^6,7,12,13^. Restricted by the sources of human RPE cells, pathology studies with human RPE are limited. In this study, we collected hRPE cells from 18 eye donors distributed over a wide age range (10 - 67 years) to analysis the molecular changes of RPE in aging process. And the cells were further confirmed by RPE-specific antibody (anti-human RPE65).

Quantitative MS-based proteome analysis with isobaric labeling reagents such as TMT have been widely used to profile the protein changes between different samples^37^. In this work, we used a MS-based isobaric relative quantitative approach for proteome and secretome analysis that provided broad coverage of the proteins of hRPE in individuals over a wide age range. We adjusted our analysis for potential confounders. The biological function of most of the proteins reported in this study was gathered by an extensive review of the literature, rather than only relying on annotation of Uniprot or GO database. We believed our approach and data were robust and could produce a descriptive quantitative dataset to show aging-associated molecular changes. An important limitation of this work is that proteomic analysis provides a static image of the protein concentration at one point in time. Therefore, we didn’t present information on dynamics of protein accumulation (or loss), or their post-translational modification (e.g. phosphorylation, acetylation). Addressing these important parameters will help us interpret more fully the biological meaning of our findings.

We described the proteome and secretome features of hRPE cells, that were not previously described. 5735 intracellular and 2455 secreted proteins were quantified in all samples (n=18), and the quantified intracellular proteins were further classified by their PANTHER protein class (version 15.0) and Gene Ontology (GO) subcellular location. RPE cells are pigment-rich cells with obvious secretory properties ^24^. In this work, we also quantified 447 secretory vesicle proteins and 81 pigment granule proteins. Besides, comparing with the expressed genes predicted by The Human Protein Atlas Database, we quantified 93 proteins that were not detected in retina and one protein not detected in any tissue.

We revealed the variances of hRPE in aging process, and uncovered the age-associated proteins and their related functions. GSEA analysis of the intracellular proteins indicated that hRPE cells was down-regulated in proteins secretion and unfolded protein response, whereas up-regulated in P53 pathway and apoptosis.

Besides, altered secretory functions like down-regulation of ER organization and up-regulation of apoptotic signaling related pathways were consist with the intracellular changes. Cells will become more vulnerable to environmental damage in aging. Endoplasmic reticulum (ER) is the site of entry of proteins into the secretory pathway ^32^. Unfolded protein response allows the cell to manage ER stress that is imposed by the secretory demands associated with environmental damage ^38^. P53 was reported to be involved in DNA repair, apoptosis and cellular stress responses, and played an important role in modulating cellular senescence and organismal aging ^39^. Therefore, molecular changes like down-regulation in protein secretion and unfolded protein response, together with up-regulation in P53 pathway and apoptosis may contribute to the pathogenesis of RPE aging.

Ubiquitin-mediated proteolysis is critical in proteostasis, which counteracts the insults accumulated to proteins by removing damaged proteins during aging. GO enrichment analysis of the age-associated intracellular proteins (e.g RNF149, RNF123, ARMC8, etc.) indicated that proteolysis involved in cellular protein catabolic process were up-regulated in aging. RNF149 and RNF123 are two E3 ubiquitin-protein ligases, which regulate protein degradation through ubiquitin-mediated proteolysis ^29^. WB results in our study showed that RNF123, RNF149 and the ubiquitinated proteins were up-regulated with the aging process of hRPE. Besides, hRPE cells from older donors were more vulnerable to oxidative damage, and they are related with the up-regulation of protein ubiquitination. Oxidative stress is a hallmark of aging and previous reports using H_2_O_2_-treated cells to analysis the mechanisms of aging ^30,31^.

Our results showed that with H_2_O_2_ treatment, RNF123, RNF149 and the ubiquitinated proteins in ARPE-19 cells were also up-regulated, and this phenomenon can be reversed by antioxidant (lutein). Therefore, RNF123 and RNF149 related protein ubiquitination may be an important clearance mechanism in elimination of oxidative damaged proteins in aged hRPE. The functions and mechanisms of RNF123 and RNF149 related protein ubiquitination during RPE aging process required more intensive studies in the following researches.

In a word, our results painted a detailed molecular picture of human RPE in aging process, and presented solid data for future research on RPE and related retinal conditions.

## ACKNOWLEDGMENTS

This work was supported by National Natural Science Foundation of China grants (82004001 and 82071008), Key Technologies Research and Development Program of Henan Science and Technology Bureau (212102310307 and 192102310076).

## DATA AVAILABILITY

The authors declare that all the other data supporting the findings of this study are available within this manuscript and its supplementary information or from the corresponding authors on request.

## CONFLICT OF INTEREST

The authors declare that there is no conflict of interest that could be perceived as prejudicing the impartiality of the research reported.

## AUTHOR CONTRIBUTIONS

X.J. designed the experiments and prepared the manuscript. J.-Y.L., W.W., X.D., and L.Y. participated in hRPE cells isolation and verification. J.-F.L. and J.-Y.L. performed MS analysis and bioinformatics analysis. G.L. and M.L. participated in secreted protein isolation. R.Q. and M.Y. helped to draft the manuscript. B.L. designed the study, provided research funding and revised the manuscript.

## Supplementary Information

### Supplementary Figures

**Supplementary Figure 1.**
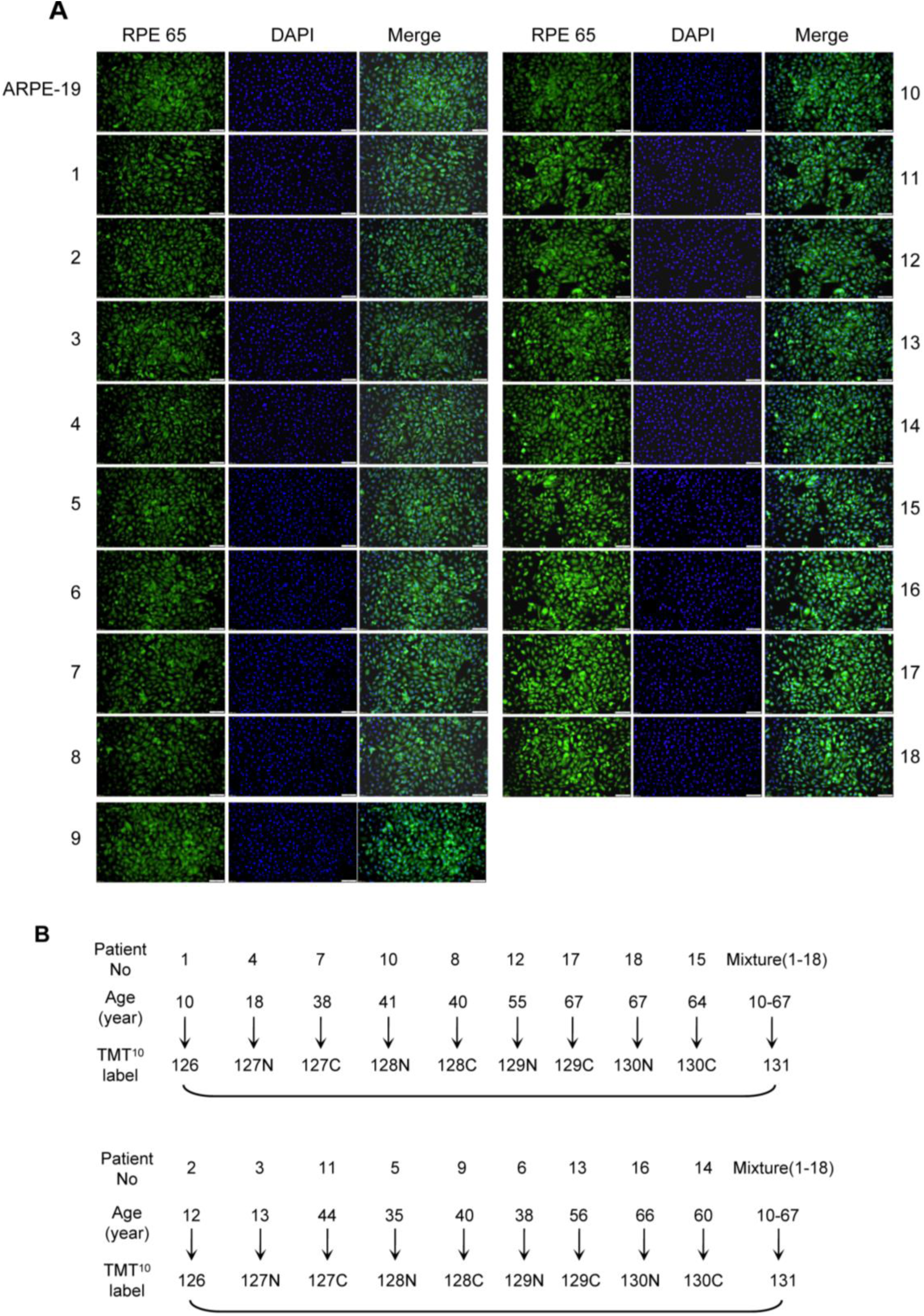
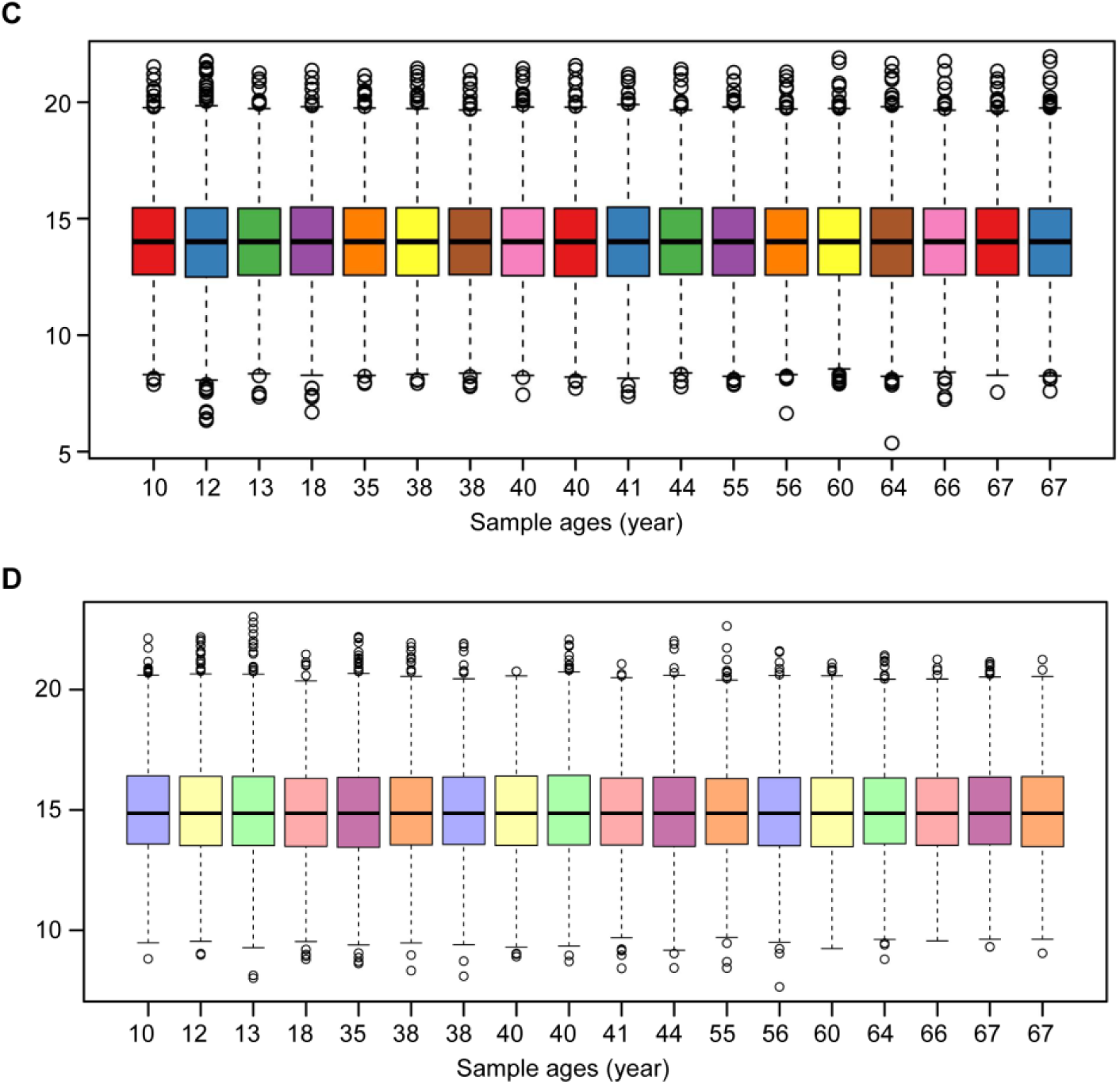
(A) Immunofluorescence staining of the hRPE cells with antibody against RPE65. RPE-specific antibody, anti-human RPE65 confirms the cell lines we collected from eyes of the donors were RPE-derived cell lines. RPE65 locates in the cytoplasm around DAPI-positive nucleus. Both hRPE and ARPE-19 cells displayed similar morphology and RPE65 staining results. (B) The detailed sample assignments for TMT^10^ labeling. Eighteen intracellular samples were randomly labeled by TMT-10plex labeling reagents, and mixture of them was labeled by TMT^131^ as internal stand in each batch of TMT-10plex. Secreted samples were labeled in the same way as intracellular sample assignments. (C-D) Boxplots indicate 75% interquartile range, median, outliers of the intracellular (C) and secreted (D) samples after a between-sample normalization by the median of intensities.

**Supplementary Figure 2.**
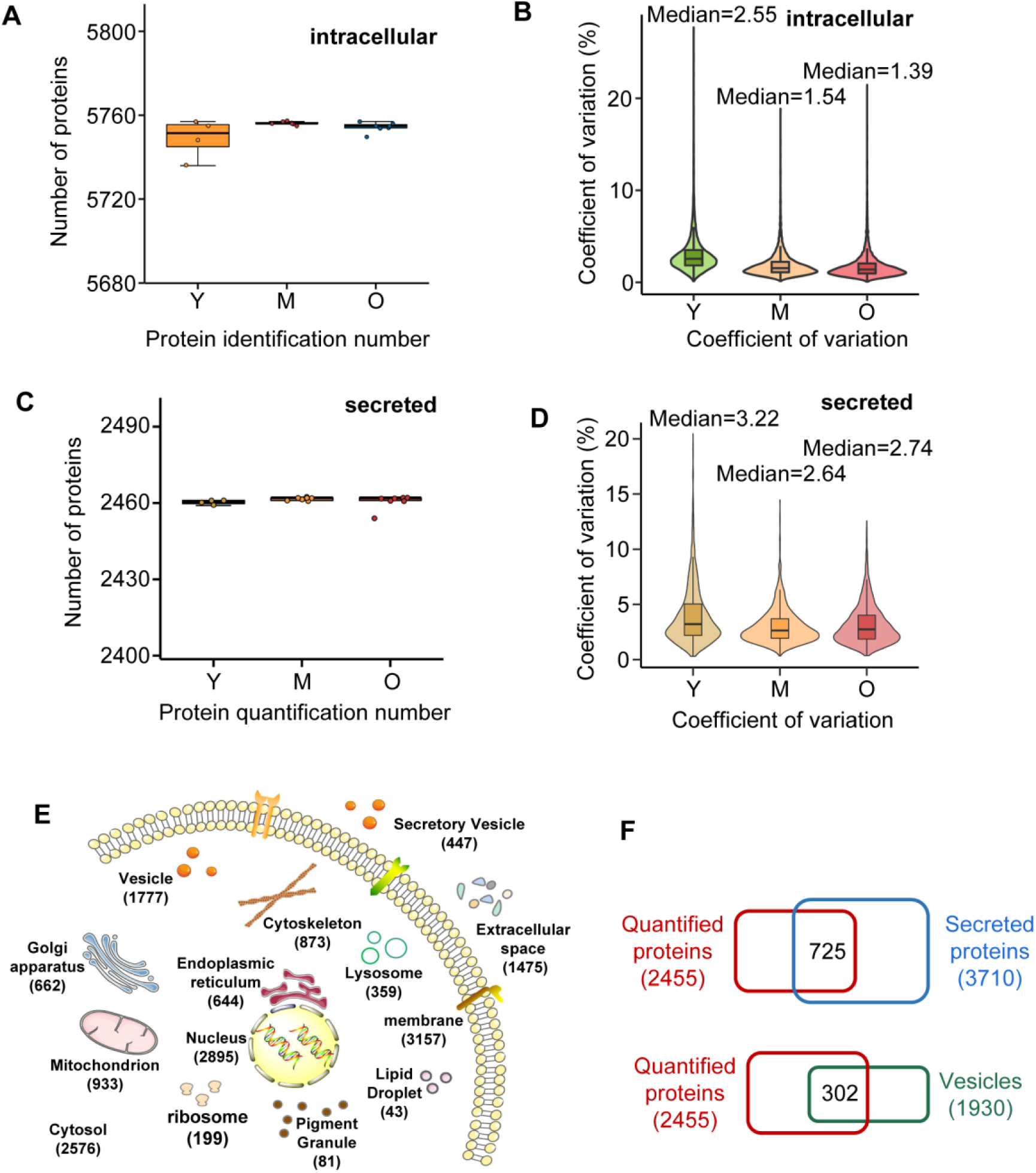
(A-D) Quality controls of the protein quantification, including number of intracellular (A) and secreted (C) proteins, and coefficient of variation (CV) of the intracellular (B) and secreted (D) proteins in the Y, M and O groups. Number of quantified proteins and the median CV values of the Y, the M and the O groups were comparable. (E) Subcellular location analysis of the intracellular proteins according to GO database. Numbers of the quantified proteins are indicated in brackets below subcellular type. (F) Venn diagram shows the overlap between secreted proteins (or vesicles) and the quantified proteins in our results. Secreted proteins or vesicles were predicated on the mRNA level according to The Human Protein Atlas Database-The Cell Atlas.

**Supplementary Figure 3.**
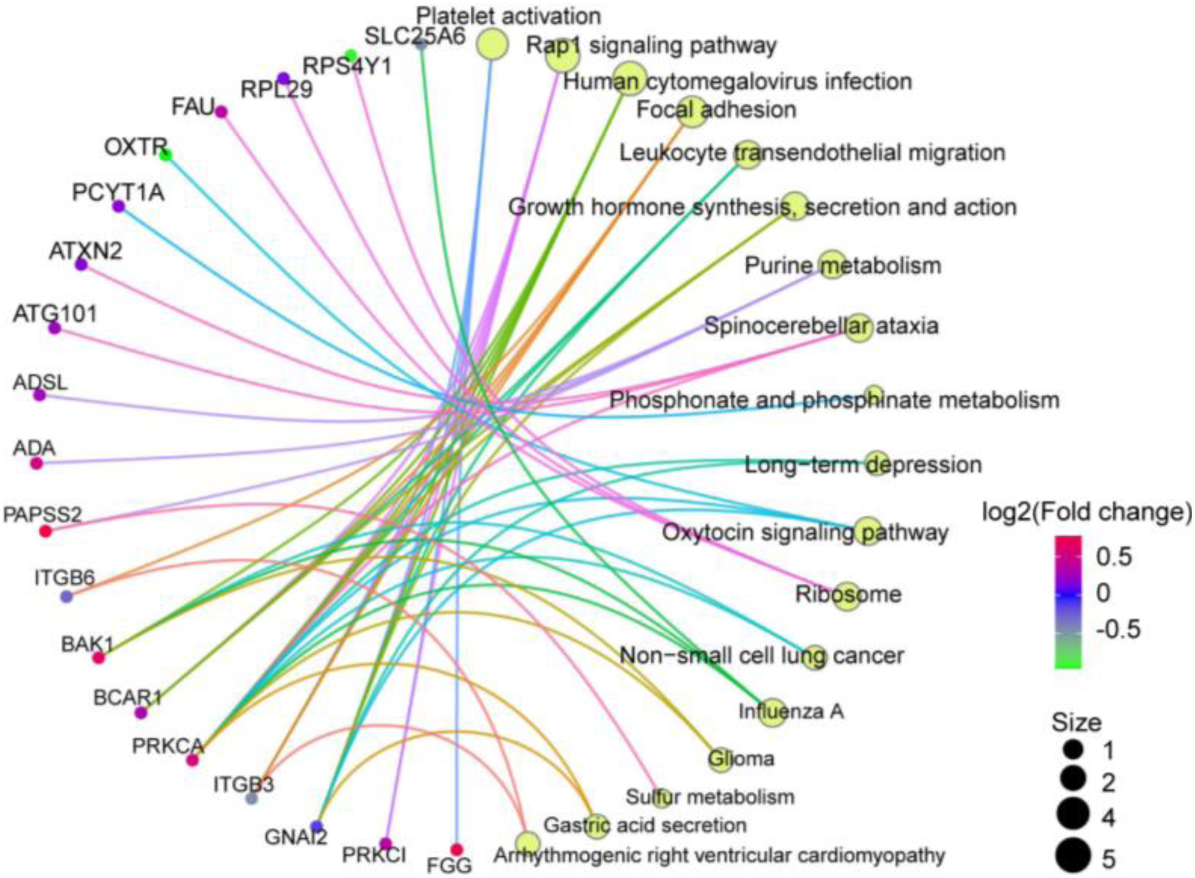
KEGG pathway analysis reveals the significantly changed (p < 0.05) pathways enriched by age-associated proteins. Dotted circle (left) denotes the proteins, circles (right) denotes their enriched pathways, connected by lines in the same color. Color of the dotted circle represents fold change. Circle size represents the number of proteins.

**Supplementary Figure 4.**
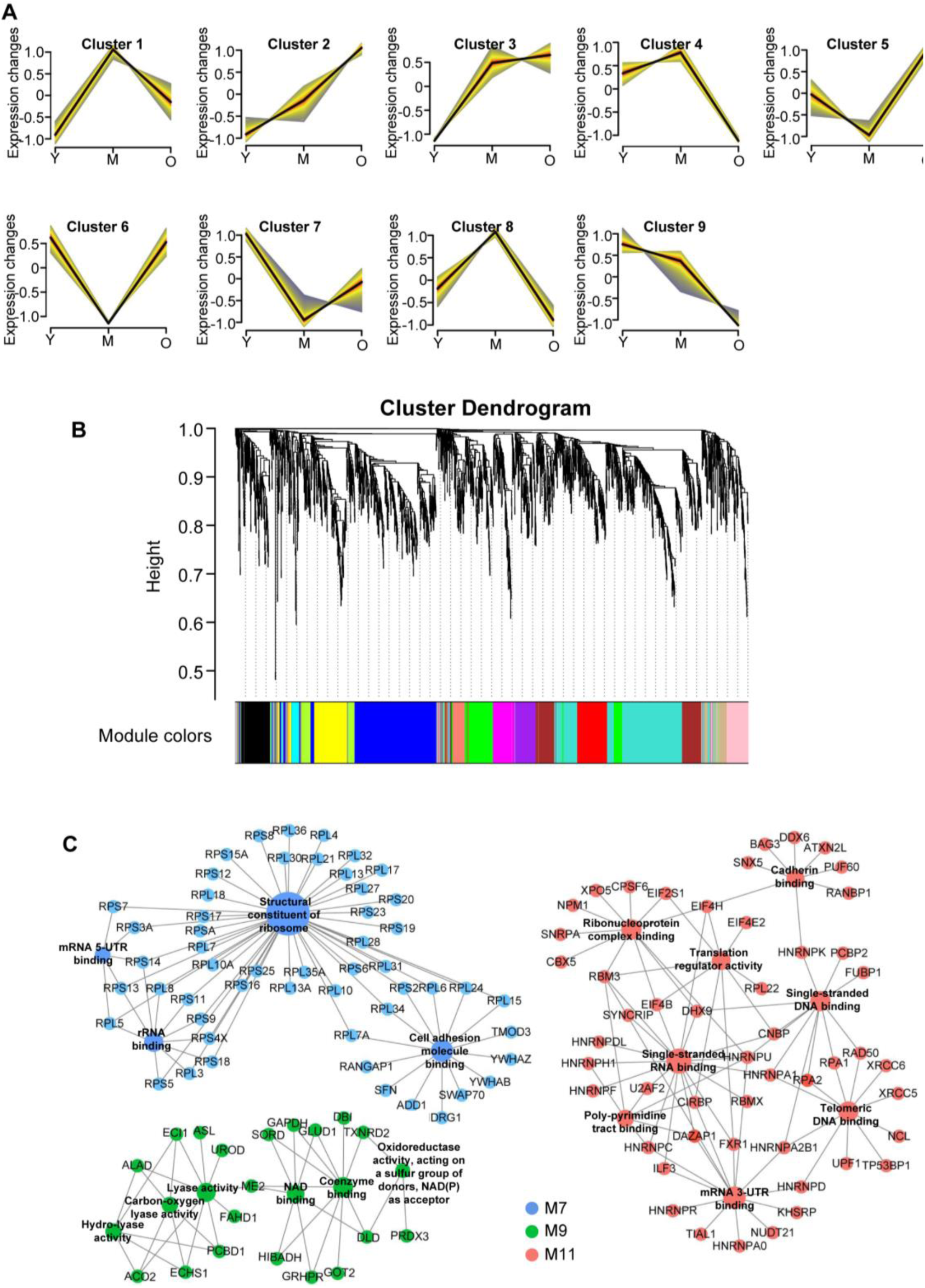
(A) Significant discrete clusters to illustrate the relative expression changes of the secreted proteomics. Clusters were clustered by mFuzz. (B) WGCNA cluster dendrogram generated by unsupervised hierarchical clustering of all proteins in the entire secreted proteomic. The analysis was based on topological overlap followed by branch cutting, and reveals 15 network modules coded by different colors. (C) GO (biological process) network depiction of protein co-expression modules. Nodes represent proteins and edges (lines) indicate connections between the nodes. Blue represents M7 (positive correlations), Green represents M9 (negative correlations), and red represents M11 (positive correlations).

### Supplementary Tables

**Supplementary Table 1.** Donor Information for hRPE cells.

| Patient Number | Age (years) | Classification | Cell Passages | Cause of Death |
| --- | --- | --- | --- | --- |
| 1 | 10 | Y | 3 | Brain death |
| 2 | 12 | Y | 3 | Accidental falling |
| 3 | 13 | Y | 3 | Accidental falling |
| 4 | 18 | Y | 3 | Craniocerebral trauma |
| 5 | 35 | M | 3 | Craniocerebral trauma |
| 6 | 38 | M | 3 | Craniocerebral trauma |
| 7 | 38 | M | 3 | Cerebral hemorrhage |
| 8 | 40 | M | 3 | Craniocerebral trauma |
| 9 | 40 | M | 3 | Cerebral hemorrhage |
| 10 | 41 | M | 3 | Craniocerebral trauma |
| 11 | 44 | M | 3 | Cerebral hemorrhage |
| 12 | 55 | O | 3 | Brain death |
| 13 | 56 | O | 3 | Cerebral hemorrhage |
| 14 | 60 | O | 3 | Cerebral hemorrhage |
| 15 | 64 | O | 3 | Acute cerebral infarction |
| 16 | 66 | O | 3 | Cerebral hemorrhage |
| 17 | 67 | O | 3 | Sudden cardiac arrest |
| 18 | 67 | O | 3 | Accidental falling |

Note: Y: The young; M: The middle aged; O: The old.

### Supplementary Data

Supplementary Data 1. Quantitative proteomic profiling of the hRPE intracellular and secreted proteins.

Supplementary Data 2. Summary of the cluster and GSEA analysis of the intracellular proteins.

Supplementary Data 3. Summary of the age-associated intracellular proteins and their functional analysis.

Supplementary Data 4. Summary of the functional analysis of the secreted proteins.

Supplementary Data 5. Summary of the specifically changed secreted proteins and their functional analysis.

## Notes

### Competing Interest Statement

The authors have declared no competing interest.

